# Optimization of the VSV G backbone for amino terminal fusion with nanobodies allowing its retargeting to receptors of therapeutic interest

**DOI:** 10.1101/2024.11.14.623577

**Authors:** Isoline Duquénois, Stéphanie Thébault, Sarah Johari, Hélène Raux, Malika Ouldali, Cécile Lagaudrière-Gesbert, Franck Perez, Aurélie A. Albertini, Yves Gaudin

## Abstract

Vesicular stomatitis virus (VSV) is a promising oncolytic virus. Additionally, its glycoprotein G is the most commonly used envelope glycoprotein to pseudotype lentiviral vectors for gene therapy. However, G receptors (LDLR family members) are ubiquitous and expressed at the surface of non-target cells, precluding *in vivo* gene therapy. Recently, we identified G mutants that no longer bind to LDLR but retain their fusion activity. This opened the possibility of specifically retargeting the glycoprotein to receptors of interest. Here, we constructed chimeric glycoproteins fused with a nanobody at the amino-terminus of G. By experimental evolution, we identified two mutations in G improving the folding and functionality of chimeric Gs, regardless of the nanobody inserted at the amino-terminus. We then constructed chimeric glycoproteins using several nanobodies targeting HER2 receptor and, into these chimeras, we introduced mutations that abolish the recognition of LDL receptors. VSV and lentiviruses pseudotyped with these glycoproteins specifically infect cells expressing HER2. We have therefore identified G mutations that optimize the G scaffold to tolerate amino-terminal insertion of a nanobody and establish proof of concept that this approach can be used to confer a new tropism on G. This paves the way for targeted *in vivo* therapies.

## Introduction

Vesicular stomatitis virus (VSV) is a promising oncolytic virus particularly for the treatment of refractory solid tumors (1–6). Indeed, it replicates preferentially in tumor cells due to their interferon (IFN) pathway defects and the subsequent lysis of infected cells induces the release of tumor antigens, which then stimulate the anti-tumor immune response (2,7). Its glycoprotein G (VSV-G) is also the most commonly used envelope glycoprotein to pseudotype lentiviral vectors for gene therapy approaches (8,9). In both cases, the lack of specificity of VSV G limits the spectrum of its use. Indeed, G binds to VSV receptors which are members of the low-density lipoprotein receptor (LDL-R) family (10) and present on the surface of almost all cell types with the notable exception of cells of the hematopoietic lineage. In virotherapy, this can lead to off-target toxicity, especially neurotoxicity (11–13). This lack of specificity also precludes *in vivo* gene therapy (*i.e.* by administering the lentiviral vector to a specific organ or area in the body). There is therefore a strong need to develop glycoproteins derived from VSV G specifically targeting cells of interest for therapeutic purposes.

VSV G has two functions essential for viral entry into the host cell (14). First, it initiates infection by binding to a cellular receptor (the LDL-R or a member of its family) which triggers endocytosis of the virion (15,16). Then, the acidic environment of the endosome induces a huge conformational change of G from its pre-fusion state toward its post-fusion state which catalyzes the fusion of the viral envelope with the endosomal membrane. This results in the release of the viral ribonucleoprotein in the cytoplasm for subsequent stages of infection.

VSV G is a type I glycoprotein, synthesized on endoplasmic reticulum-bound ribosomes. After cleavage of its amino terminal signal peptide, the mature glycoprotein is 495 aa long. It is anchored in the membrane by a single α-helical transmembrane (TM) segment (Figure 1A). Most of the mass of G (residues 1-446) is located outside the viral membrane and constitutes its amino-terminal ectodomain. The structures of pre- and post-fusion trimers of VSV G were determined revealing the details of the conformational change (17–19). More recently, crystal structures of the prefusion conformation of G ectodomain in complex with two cysteine-rich domains (CR2 and CR3) of the LDL-R of G have also been determined (20). The binding site of the CR domains on G was the same in both crystals and allows the identification of two basic residues (K47 and R354) which are essential for the interaction between G and its receptor. Mutant glycoproteins in which those two residues are replaced by either an alanine or a glutamine are unable to bind LDL-R CR domains and cannot pseudotype a recombinant VSV (VSVΔG-GFP) in which the G gene has been replaced by the green fluorescent protein (GFP) gene (20). Nevertheless, these mutant glycoproteins retain their ability to induce syncytia formation at low pH when expressed in cells by plasmid transfection (20). Therefore the fusion activity and receptor binding activity of VSV G can be decoupled which paves the way for the development of G-derived glycoproteins with modified tropism.

**Figure 1:**
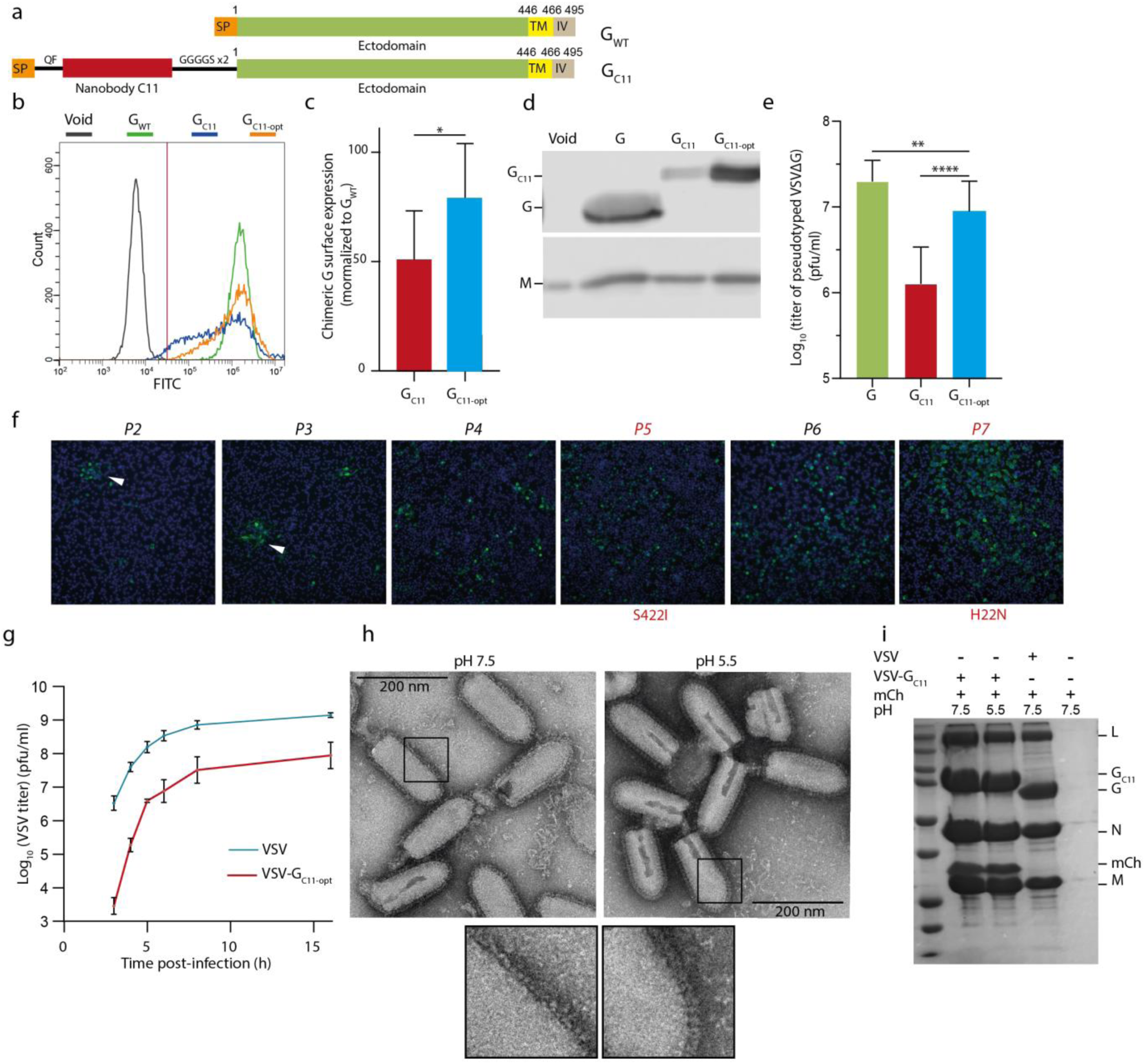
Optimisation of the VSV G backbone to tolerate an aminoterminal fusion with the C11nanobody. **A)** Schematic representation of G_WT_ and of the G_C11_ chimera. SP: signal Peptide; TM: transmembrane domain; IV: intraviral domain. **B)** Presence of G_WT_, G_C11_, and G_C11-opt_ at the cell surface of transfected HEK293T cells. G was detected on the surface of non-permeabilized cells using mAb 8G5F11 and a secondary antibody conjugated to Alexa Fluor 488. Cells were analyzed by flow cytometry. **C)** Median surface concentration of G_C11_ and G_C11-opt_ normalised to median concentration of G_WT_ on the surface of HEK293T cells. The median fluorescence was measured using flow cytometry. **D)** Incorporation of G_WT_, G_C11_, and G_C11-opt_ into VSVΔG-GFP viral particles assessed by western blot using an anti-VSV G and an anti-VSV M antibody. **E)** Infectivity of VSVΔG-GFP pseudotyped with G_WT_, G_C11_ and G_C11-opt_ in HEK293T cells. VSVΔG-GFP viruses pseudotyped with WT or chimeric glycoproteins, produced in the same conditions (see material and methods), were used to infect indicated cell lines during 16 h. The percentage of infected cells was measured by counting GFP expressing cells by flow cytometry and used to calculate the viral titer. **F)** Experimental evolution of rVSV-G_C11_ recombinant virus in BSR cells. rVSV-G_C11_ was serially passaged in BSR cells. Infected cells were labelled using an anti-N antibody and a goat anti-mouse IgG secondary antibody conjugated to Alexa Fluor 488. DAPI was used to stain nuclei. Genomic RNA was extracted and sequenced after passage 5, 7 and 10 revealing the invasion of G mutations H22N (after P5) and S422I (after P7) into the viral population. **G)** Growth curves of VSV_WT_ and rVSV-G_C11-opt_. At each time point, the viral titer was measured using plaque assay. **H)** Negative staining electron microcopy of rVSV-G_C11-opt_ incubated at pH 7.5 and 5.5. Insets show the magnification of the viral membrane revealing the typical shape of VSV G at high and low pH. **I)** Binding of mCherry protein to rVSV-G_C11-opt_. Indicated viruses were incubated with mCherry at indicated pHs, centrifuged and the pellet was analysed by SDS-PAGE. L, N, M indicate the position of viral proteins, G and G_C11-opt_ the G proteins and mCh the precipitated mCherry. In **C)**, **E)**, and **G)**, data points represent replicates of at least three independent experiments. Errors bars indicate SD. In **C)** and **E),** statistically significant differences between G, G_C11_ and G_C11-opt_ are indicated by stars (*p < 0.05, **p < 0.005, ****p < 0.00005).

A possible strategy to retarget the VSV glycoprotein toward a receptor of interest expressed at the cell surface consists of fusing it to a protein domain specifically binding this receptor. Nanobodies (also known as VHH or single domain antibodies) are an interesting class of specific binders (21,22). They are stable, rather small, and several libraries that are available for screening of high affinity bindings against a specific target (23,24). Thus, we decided to construct chimeric glycoproteins with a nanobody inserted between the signal peptide (SP) and the ectodomain of G. However, we observed that such a fusion of a nanobody (directed against mCherry) at the N-terminus of G affected the properties of the glycoprotein. We thus decided to carry out experimental evolution to optimize the backbone of G so that it tolerates this insertion. Our hypothesis was that selecting mutations that optimize the G backbone for insertion of a given nanobody should also optimize it for the insertion of any nanobody due to their common 3D structure. For this, we constructed a recombinant VSV in which the gene encoding G was replaced by a gene encoding the chimeric glycoprotein. The recombinant virus had a very low titer and after a few passages, two mutations were selected in the G backbone that were shown to optimize several chimeras involving distinct nanobodies (as assessed in VSVΔG-GFP pseudotyping experiments). We then constructed several chimeras in which this optimized G backbone was fused to distinct nanobodies directed against the receptor tyrosine-protein kinase erbB-2 (HER2, overexpressed at the surface of malignant cells (25) and used for targeted drug delivery (26)) and introduced the mutations K47Q or R354Q abolishing the recognition of LDL-R by G (20). We demonstrated that these chimeras efficiently pseudotype both VSVΔG-GFP and lentiviruses and specifically retarget them toward HER2-expressing cells. This work therefore provides a VSV-G backbone that makes it possible to construct chimeric Gs specifically targeting cells of interest using nanobodies.

## Results

### Construction and characterization of a chimeric glycoprotein fused to a nanobody

We constructed a chimeric glycoprotein (G_C11_) with a nanobody (C11, directed against mCherry) (24) inserted between the signal peptide (SP) and the ectodomain so that the nanobody will be located at the amino-terminal end of the mature G. The nanobody was flanked by an insertion of a QF dipeptide (to optimize the SP cleavage) and a flexible linker (GGGGSGGGGS) before G ectodomain (Figure 1A).

We transfected HEK293T cells with the resulting construct to assess G_C11_ surface expression. Non-permeabilized cells were labeled using the monoclonal antibody (mAb) 8G5F11, directed against G ectodomain and quantified by flow cytometry. Surface expression of G_C11_ was affected compared to that of the wild type glycoprotein (G_WT_) (Figure 1B) and the median density of G_C11_ protein present at the cell surface was 50 +/- 21% of that of G_WT_ (Figure 1C).

We also assessed the ability of G_C11_ to pseudotype a recombinant VSV (VSVΔG-GFP) in which the G envelope gene was replaced by the green fluorescent protein (GFP) gene. Chimeric glycoproteins incorporation into the envelope of the particles present in the supernatant was verified by western blot. Incorporation of the G_C11_ chimera into the viral membrane was less efficient than that of G_WT_ (Figure 1D). Pseudotyped viruses were then used to infect HEK293T cells and their infectivity was analyzed by counting GFP-expressing cells using flow cytometry. The infectious titer of the production of VSVΔG-GFP pseudotyped with G_C11_was 1.3 Log_10_ (*i.e.* 20 times) lower that of VSVΔG-GFP pseudotyped with G_WT_ (Figure 1E). This indicated that the fusion of the nanobody at the N-terminus of G affected the properties of the glycoprotein.

### Optimization of G backbone by experimental evolution

We decided to perform experimental evolution to optimize the backbone of G so that it tolerates an N-terminal fusion with a nanobody. For this, we introduced the gene encoding G_C11_ in place of that encoding G in the viral genome to construct the rVSV-G_C11_ recombinant virus. The use of nanobody C11, which recognizes mCherry, in this construct was a carefully considered choice to avoid uncontrolled gain of function (GOF). We hypothesized that selection for mutations optimizing the G backbone for insertion of the C11 nanobody would also optimize the G backbone for insertion of other nanobodies. The resulting recombinant virus only formed small infection foci in the plaque assay (Figure 1F) and has a titer of ∼10^5^ pfu/ml (Table 1). We decided to serially passage the recombinant virus to select mutants with a better fitness. After 4 passages, mutation S422I (Table 1) has invaded the viral genome population with only minor changes in the infectious phenotype. After 7 passages, we observed a strong increase in the viral infectivity. This was due to the selection of a second mutation H22N (Table 1). No other mutation was detected after 10 passages and we decided to stop to passage the virus (Figure 1F). Of note, mutation S422I had been selected in a different context (as a compensatory mutation of the deleterious mutation H407A) and demonstrated to stabilize the prefusion state (19).

**Table 1:** Adaptation of rVSV-G_C11_. Passage number with the corresponding titer on BSR cells and the identified mutation in G ectodomain. The original codon and the codon in G_C11-opt_ are indicated

| Passage | Viral titer | Mutation | Original codon | G<sub>C11-opt</sub> codon |
| --- | --- | --- | --- | --- |
| P1 | 8.5 10<sup>4</sup> pfu/ml |  |  |  |
| P2 | 1.5 10<sup>5</sup> pfu/ml |  |  |  |
| P3 | 1.4 10<sup>6</sup> pfu/ml |  |  |  |
| P4 | 1.1 10<sup>6</sup> pfu/ml | H22N | CAT | AAT |
| P5 | 2.9 10<sup>6</sup> pfu/ml |  |  |  |
| P6 | 2 10<sup>6</sup> pfu/ml |  |  |  |
| P7 | 7 10<sup>7</sup> pfu/ml | S422I | AGT | ATT |
| P8 | 7 10<sup>7</sup> pfu/ml |  |  |  |
| P9 | 9 10<sup>7</sup> pfu/ml |  |  |  |

### Characterization of recombinant rVSV-G_C11-opt_ and its glycoprotein

We characterized the resulting virus rVSV-G_C11-opt_ (opt for optimized indicating that G contains mutations H22N and S422I). We compared the one-step growth curve of rVSV-G_C11-opt_ with that of VSV WT. The titer of the recombinant virus after 16h of infection of BSR cells was ∼10^8^pfu/ml i.e. one log lower than that of VSV WT (Figure 1G).

We also investigated the structure of G_C11-opt_ at the surface of the recombinant virus using electron microscopy after negative staining. At pH 7.5, both G_WT_ and G_C11-opt_ formed the characteristic ∼8-nm-wide layer (27). At pH 5.5, at the surface of the viral particles, the spikes were individualized, allowing the visualization of 12-nm-long spikes in their post-fusion conformation that stood perpendicular to the membrane (Figure 1H) (27). This indicated that G_C11-opt_ undergoes the low pH-induced structural transition.

We then verified the ability of G_C11-opt_ at the surface of the recombinant virus to bind to purified mCherry at pH 7.5 and 5.5. For this, the virus was incubated with mCherry at indicated pH before ultracentrifugation through a sucrose cushion. Under both pH conditions, mCherry was found associated with the recombinant virus but not with the WT virus in the pellet indicating G_C11-opt_ binds the mCherry via the C11 nanobody (Figure 1I). The C11 nanobody is thus folded and accessible in both the pre- and post-fusion states of G at the viral surface.

We also compared the surface expression of G_C11-opt_ with that of G_WT_ and G_C11_. Cells were transfected with plasmids encoding the corresponding glycoproteins and the surface expression was quantified by flow cytometry using mAb 8G5F11. G_C11-opt_ was significantly more expressed at the cell surface than G_C11_. Indeed, the median density of G_C11-opt_ protein present at the cell surface was 79 +/- 23% of that of G_WT_ (Figure 1B, 1C). Therefore, mutations H22N and S422I improve G transport to the surface.

Finally, we compared the ability of G_C11-opt_, G_WT_, and G_C11_ to pseudotype VSVΔG-GFP. Chimeric glycoproteins incorporation into the envelope of the viral particles was verified by western blot revealing that G_C11-opt_ chimera was more efficiently incorporated than G_C11_ (Figure 1D). Pseudotyped viruses were then used to infect HEK293T cells and their infectivity was analyzed by counting GFP-expressing cells using flow cytometry (Figure 1E). The infectious titer of the production of VSVΔG-GFP pseudotyped with G_C11-opt_ (∼10^6.95^ pfu/ml) was slightly lower that of VSVΔG-GFP pseudotyped with G_WT_ (∼10^7.3^ pfu/ml) and ∼0.85 log_10_ (*i.e.* 7 fold) higher than that of the VSVΔG-GFP pseudotyped with G_C11_ confirming that the two mutations H22N and S422I optimize the infectivity of the chimera.

### Hydrophobic residues at position 422 combined with Asn at position 22 optimize G for nanobody insertion

The structure of VSV G ectodomain in its pre-fusion conformation reveals that residue S422 points toward F424 and L430. The selection of an isoleucine in position 422 probably stabilizes the β-hairpin structure through hydrophobic stacking of residues I422 between F424 and L430 (Figure 2A). The codon encoding S422 in the G_WT_ gene is AGU (Table 1) and the only codon encoding a hydrophobic residue that is accessible by a single nucleotide change is AUU encoding the isoleucine. Consequently, other hydrophobic amino acids were not directly accessible during experimental evolution. We decided to test the impact of other hydrophobic amino acid residues (L, V, F, M) in this position. Thus, we replaced isoleucine 422 in G_C11-opt_ by a leucine, a valine, a phenylalanine, or a methionine leading to constructs G_C11-opt-L422_, G_C11-opt-V422_, G_C11-opt-F422_, and G_C11-opt-M422_ (note that all these constructs contain the H22N mutation). We then analyzed the ability of these constructs to pseudotype VSVΔG-GFP. The infectious titers of VSVΔG-GFP pseudotyped with chimeric glycoproteins G_C11-opt-L422_, G_C11-opt-V422_, G_C11-opt-F422_, or G_C11-opt-M422_ were similar to that of VSVΔG-GFP pseudotyped with G_C11-opt_, significantly higher that of VSVΔG-GFP pseudotyped with G_C11_, but not significantly different from that of VSVΔG-GFP pseudotyped with G (Figure 2B). This demonstrated the importance of a hydrophobic residue at position 422 for optimizing G backbone for nanobody insertion.

**Figure 2:**
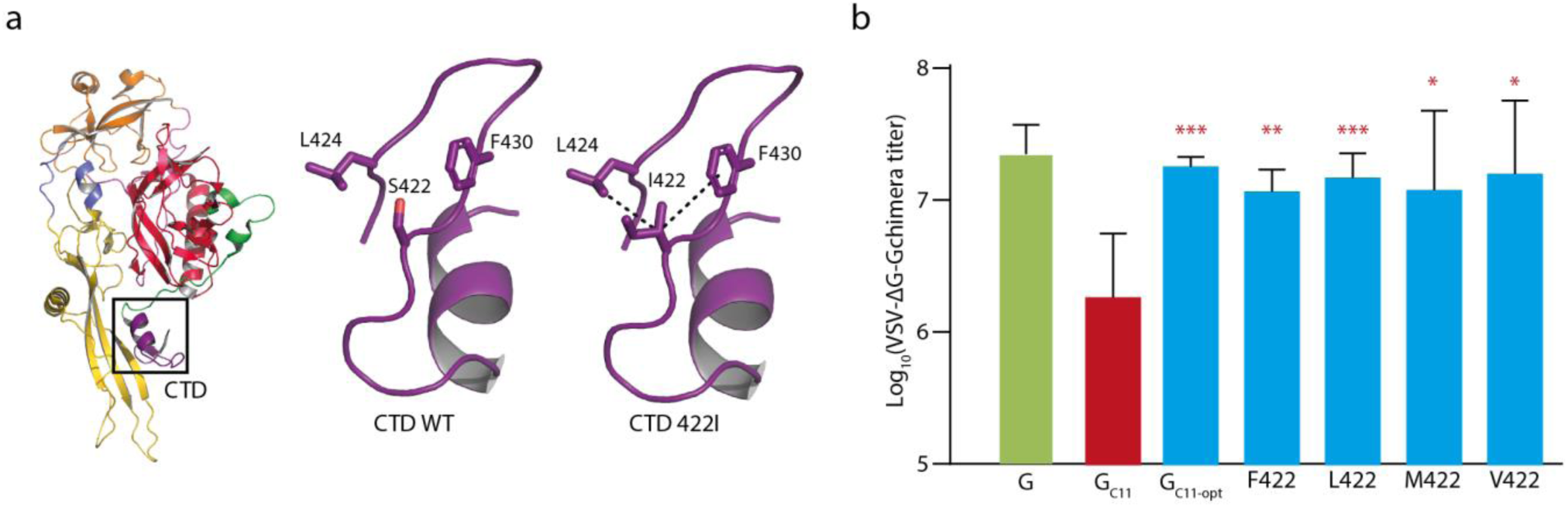
Hydrophobic residues at position 422 optimize G for nanobody insertion. **A)** Environment of residue S422 in the crystal structure of the prefusion conformation of VSV G. **B)** Mutation S422I stabilizes G C-terminal domain (CTD) through hydrophobic stacking with residues F424 and L430. **C)** Infectivity of VSVΔG-GFP pseudotyped with G, G_C11_, _GC11-opt_, G_C11-opt-S422F_, G_C11-opt-S422L_, G_C11-opt-S422M_, and G_C11-opt-S422V_ in HEK293T cells. VSVΔG-GFP viruses pseudotyped with G_WT_ or chimeras, produced in the same conditions (see material and methods), were used to infect HEK293T during 16 h. The percentage of infected cells was measured by counting GFP expressing cells by flow cytometry and used to calculate the viral titer. Data points represent replicates of five independent experiments. Errors bars indicate SD. Statistically significant differences with G_C11_ (resp. G) are indicated by red (resp. green) asterisks (*p < 0.05, **p < 0.005, ***p < 0.0005, nxs not significant).

### The optimized G backbone is functional for other nanobodies insertion

We investigated whether the double mutation H22N and S422I also improved the functionality of G chimeras with other nanobodies. For this, we inserted in the G_WT_ or G_opt_ backbones 4 other nanobodies from the same library (24): 8G5 directed against the carcinoembryonic antigen cell adhesion molecule (CEACAM), 9C2 directed against mucin 1 protein (MUC1), C8 and A10 both targeting HER2 and 6 others (H1 to H6) also targeting HER2, whose sequences were available in databanks.

The resulting constructions were called G_C8_, G_8G5_, G_A10_, G_9C2_, G_H1_ G_H2_ G_H3_ G_H4_, G_H5,_ and G_H6_ when the insertion was in the backbone of G_WT_, and G_C8-opt_, G_8G5-opt_, G_A10-opt_, G_9C2-opt_, G_H1-opt_, G_H2-opt_, G_H3-opt_, G_H4-opt_, G_H5-opt_, and G_H6-opt_ when the insertion was in the optimized G backbone containing the mutations H22N and S422I.

For each nanobody, we then compared the ability of the corresponding pair of chimeras (*i.e.* optimized or non-optimized) to pseudotype VSVΔG-GFP. For this, HEK293T cells were infected by the viruses pseudotyped with the chimeras. For 9 nanobodies out of 11 (including C11), the infectivity in HEK293T cells of pseudotype production was enhanced when the nanobody was inserted in the optimized backbone. The enhancement factor varied from 6.5 to 29 depending on the nanobody. The geometric mean of the increase for these nine nanobodies was 11 (Figure 3A and table 2). For two chimeras (G_8G5_ and G_9C2_), the optimized backbone had no significant effect on the infectivity of the chimera.

**Figure 3:**
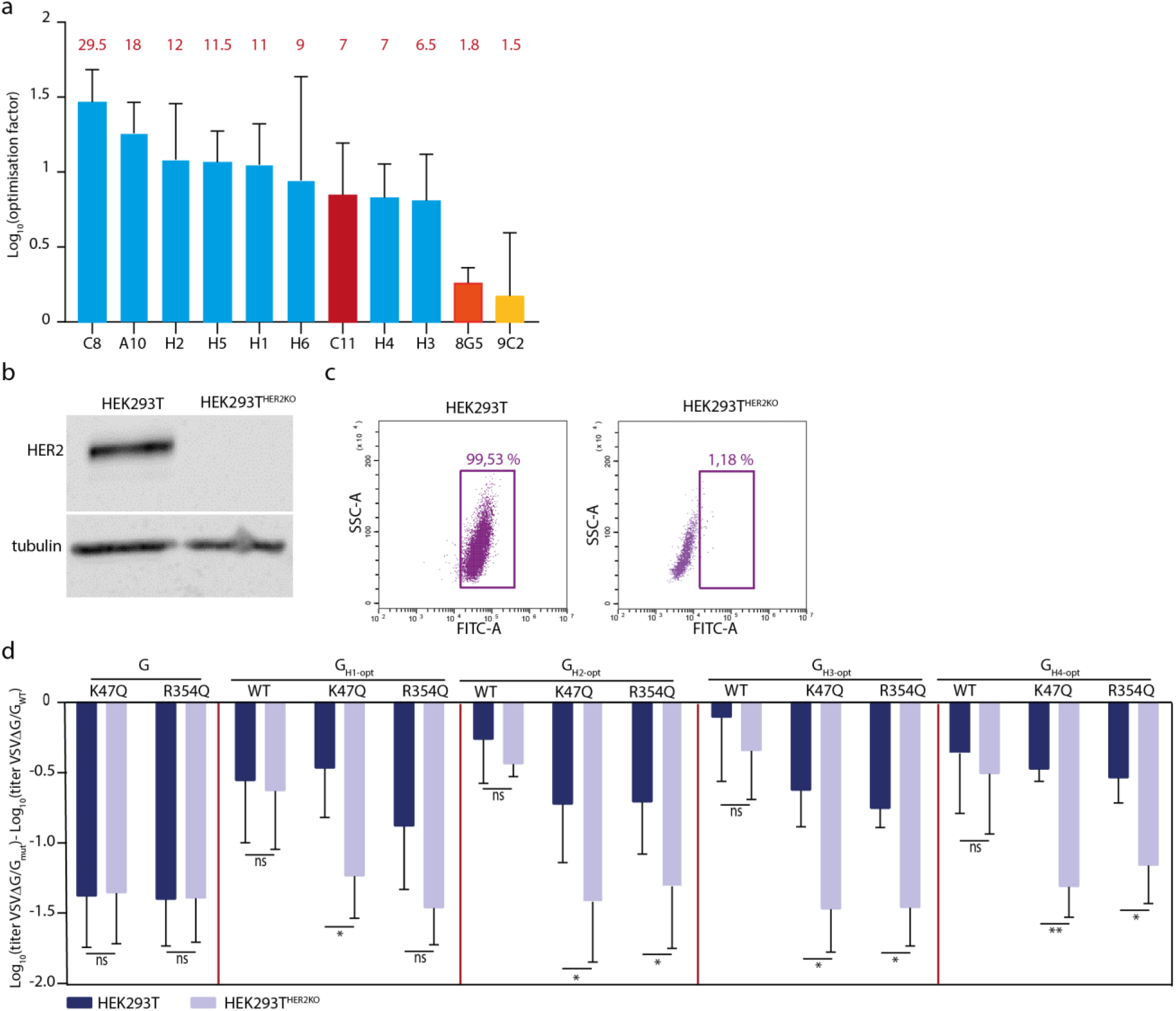
Retargeting of VSVΔG-GFP towards HER2-expressing cells. **A)** Optimization factor of the titer of VSVΔG-GFP pseudotyped with the chimeric G in the optimized backbone compared with VSVΔG-GFP pseudotyped with a chimera in a non-optimized backbone. VSVΔG-GFP viruses, produced in the same conditions (see material and methods) pseudotyped with chimeras in an optimized or non-optimized backbone were used to infect indicated cell lines during 16 h. The percentage of infected cells was measured by counting GFP expressing cells by flow cytometry and used to calculate the viral titer and used to calculate the viral titer. The histogram shows the mean difference between the Log_10_ of the titer of VSVΔG-GFP pseudotyped with the optimized chimera and the Log_10_ of the titer of VSVΔG-GFP pseudotyped with the non-optimized chimera. The factor of optimization is indicated in red for each chimera. **B)** Expression of HER2 in HEK293T and HEK293T-HER2^KO^ cells. Cell lysates were analysed by western blot with anti-HER2 and anti-tubulin antibodies. **C)** Cell surface expression of HER2 in in HEK293T and HEK293T-HER2KO cells. Expression of HER2 was analysed using flow cytometry with an anti-HER2 as a primary antibody and a secondary antibody conjugated to Alexa Fluor 488. The percentage of HER2-positive cells is indicated. **D)** Infectivity in HEK293T and HEK293T-HER2^KO^ cells of VSVΔG-GFP pseudotyped with G_K47Q_, G_R354Q_ or by optimized G chimeras fused to nanobodies H1 to H4 (recognizing HER2) in which mutations K47Q or R354Q have been introduced or not (WT). VSVΔG-GFP viruses pseudotyped with these glycoproteins were used to infect indicated cell lines during 16 h. The percentage of infected cells was measured by counting GFP expressing cells by flow cytometry and used to calculate the viral titer. The histogram indicates the Log10 of the titer of viruses pseudotyped with mutated or chimeric Gs normalized to that pseudotyped with G_WT_ in the indicated cell line. The Log_10_ of the titer of VSVΔG-GFP virus pseudotyped with G_WT_ was 7.37 +/- 0.34 in HEK293T and 7.39 +/- 0.38 in HEK293T-HER2^KO^ cells (n=13). In **A)** and **D),** error bars represent the SD for experiments carried out in at least in triplicate (see also table 2 for A). All experiments were at least performed in triplicate. Statistically significant differences are indicated by stars (*p < 0.05, **p < 0.005, ns: not significant).

**Table 2:** Optimization factor for each chimera (indicated by its fused nanobody) in the optimized G backbone context versus the non-optimized G context in HEK293T cells. The titer of VSVΔG-GFP pseudotyped with the optimized chimera is also indicated as well as the nanobody target and the number of experiments performed. The Log_10_ of the titer of VSVΔG-GFP pseudotyped with G_WT_ (used as a reference in each experiment) was 7.3 +/- 0.25 (n=16).

| Nanobody | Log<sub>10</sub>(optimization factor) +/- SEM | Optimization factor | Log<sub>10</sub>(Titer of optimized pseudotype) | Target | n |
| --- | --- | --- | --- | --- | --- |
| C11 | 0.85 +/- 0.35 | 7 | 6.95 +/- 0.35 | mCherry | 16 |
| C8 | 1,47 +/- 0.21 | 29,5 | 6.34 +/- 0.3 | HER2 | 3 |
| A10 | 1.26 +/- 0.21 | 18 | 6.08 +/- 0.27 | HER2 | 3 |
| H1 | 1.05 +/- 0.27 | 11 | 6.95 +/- 0.16 | HER2 | 4 |
| H2 | 1.08 +/- 0.38 | 12 | 6.75 +/- 0.19 | HER2 | 5 |
| H3 | 0.81 +/- 0.31 | 6,5 | 7.24 +/- 0.48 | HER2 | 4 |
| H4 | 0.83 +/- 0.22 | 7 | 7.23 +/- 0.48 | HER2 | 4 |
| H5 | 1.07 +/- 0.21 | 11,5 | 6.59 +/- 0.23 | HER2 | 5 |
| H6 | 0.94 +/- 0.69 | 9 | 6.71 +/- 0.46 | HER2 | 3 |
| 8G5 | 0,26 +/- 0.1 | 1.8 | 4.94 +/- 0.31 | CEACAM | 3 |
| 9C2 | 0.17 +/- 0.42 | 1.5 | 5.18 +/- 0.43 | Muc1 | 3 |

### Retargeting of pseudotyped VSVΔG-GFP toward cells expressing HER2 receptor

We showed that the optimization mutations that we had selected with the G_C11_ chimera also improved most of the chimeras involving other nanobodies. However, all of these chimeras remain capable of recognizing the natural VSV receptor (LDL-R).

We therefore introduced the mutations K47Q or R354Q, abolishing receptor recognition by VSV G (20), in the optimized chimera in which G is fused with nanobodies directed against HER2. We chose to work with chimeras involving nanobodies H1 to H4 because these chimeras had the best infectious titers in their optimized version (Table 2). We thus constructed G_H1-opt-K47Q_, G_H1-opt-R354Q_, G_H2-opt-K47Q_, G_H2-opt-R354Q_, G_H3-opt-K47Q_, G_H3-opt-R354Q_, G_H4-opt-K47Q_, and G_H4-opt-R354Q_.

As HER2 was expressed at the surface of HEK293T (Figure 3B and 3C), we constructed a HEK293T-HER2^KO^ cell line in which the *ERBB2* gene (encoding HER2 protein) was knocked out (Figure 3B and 3C). We compared the infectivity of VSVΔG-GFP pseudotyped with G_WT_, G mutants incapable of recognizing LDL-R (G_K47Q_ and G_R354Q_) and the different chimeras involving nanobodies H1 to H4 (capable or not of recognizing LDL-R). As expected, the titer of VSVΔG-GFP pseudotyped with G_K47Q_, G_R354Q_, was reduced by a factor of ∼25 (1.4 Log_10_) in both cell lines compared to G_WT_ (Figure 3D). The chimeric glycoproteins G_H1-opt_, G_H2-opt_, G_H3-opt_, G_H4-opt_, still able to recognize LDL-R, efficiently pseudotyped VSVΔG-GFP in both cell lines, nevertheless somewhat less efficiently than G_WT_ in agreement with table 2. Finally, the titer of VSVΔG-GFP pseudotyped with G_H1-opt-K47Q_, G_H1-opt-R354Q_, G_H2-opt-K47Q_, G_H2-opt-R354Q_, G_H3-opt-K47Q_, G_H3-opt-R354Q_, G_H4-opt-K47Q_, or G_H4-opt-R354Q_ was only slightly reduced in HEK293T compared to G_WT_ cells and similar to that of G_K47Q_, G_R354Q_ in HEK293T-HER2^KO^ cells. This indicated that the entry in HEK293T cells of VSVΔG-GFP pseudotyped with the different chimeras involving nanobodies H1 to H4 and unable to recognize LDL-R is mediated by HER2 (Figure 3D).

### Chimeras are also functional to retarget lentiviral vectors

VSV G is the most used viral glycoprotein to pseudotype lentivirus vectors. We decided to investigate if the chimera in which G, carrying the mutation K47Q or R354Q, is fused with nanobodies directed against HER2 could specifically redirect lentiviruses toward HER2-expressing cells. To do this, we compared the infectivity of GFP-encoding lentivirus pseudotyped with the different chimeras in HEK293T and HEK293T-HER2^KO^. Infectivity was analyzed by counting GFP-expressing cells using flow cytometry (Figure 4A).

**Figure 4:**
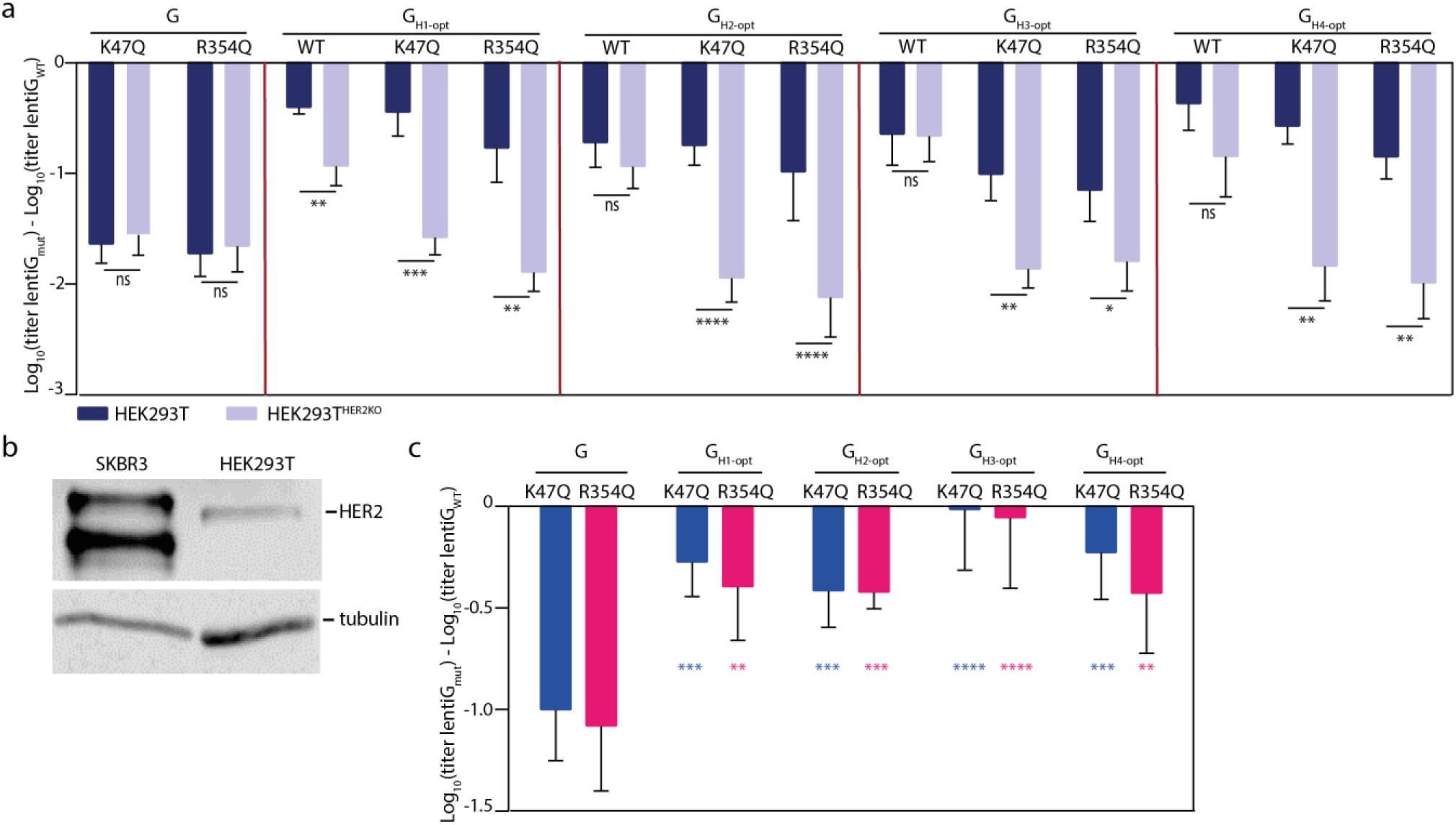
Infectivity of lentiviruses pseudotyped with G_K47Q_, G_R354Q_ or by optimized G chimeras fused to nanobodies H1 to H4 (recognizing HER2) in which mutations K47Q or R354Q have been introduced or not (WT). **A)** Comparison of infectivity of lentiviruses pseudotypes by the indicated G construct in HEK293T and HEK293T-HER2^KO^ cells. The percentage of transduced cells was measured by counting GFP expressing cells by flow cytometry 48h after transduction. The histogram indicates the Log_10_ of the titer of lentiviruses pseudotyped with the indicated G construct normalised with the one pseudotyped with G_WT_. The Log_10_ of the titer of lentiviruses pseudotyped with G_WT_ was 6.89 +/- 0.22 in HEK293T and 6.83 +/- 0.22 (n=13) in HEK293T-HER2^KO^ cells. **B)** Comparison of the expression of HER2 in HEK293T and SKBR3. Cell lysates were analysed by western blot with anti-HER2 and anti-tubulin antibodies. **C)** Infectivity of lentiviruses pseudotyped with G_K47Q_, G_R354Q_ or by optimized G chimeras fused to nanobodies H1 to H4 (recognizing HER2) in which mutations K47Q or R354Q have been introduced in SKBR3 cells. The percentage of transduced cells was measured by counting GFP expressing cells by flow cytometry 48h after transduction. The histogram indicates the Log_10_ of the titer of lentiviruses pseudotyped with the indicated G construct normalised with the one pseudotyped with G_WT_. The Log_10_ of the titer of lentiviruses pseudotyped with G_WT_ was 5.8 +/-0.15 (n=10) in SKBR3 cells. In **A)** and **C),** error bars represent the SD for experiments carried out in at least in triplicate. Statistically significant differences are indicated by stars (*p < 0.05, **p < 0.005, ***p < 0.0005, ****p < 0.00005, ns: not significant). In C), blue (resp. pink) stars indicate the significance of the difference between G_K47Q_ (resp. GR354Q) and the chimera having the same mutation.

The results indicated that lentivirus pseudotyped with chimeras G_H1-opt-K47Q_, G_H1-opt-R354Q_, G_H2-opt-K47Q_, G_H2-opt-R354Q_, G_H3-opt-K47Q_, G_H3-opt-R354Q_, G_H4-opt-K47Q_, or G_H4-opt-R354Q_ infected much more efficiently HEK293T that HEK293T-HER2^KO^ (Figure 4A). This indicated that the lentiviruses were retargeted against cells that express HER2.

This was confirmed using SKBR3 cells that express high level of HER2 at their surface (Figure 4B). Lentiviruses pseudotyped with chimeras G_H1-opt-K47Q_, G_H1-opt-R354Q_, G_H2-opt-K47Q_, G_H2-opt-R354Q_, G_H3-opt-K47Q_, G_H3-opt-R354Q_, G_H4-opt-K47Q_, or G_H4-opt-R354Q_ infected more efficiently this cell line than those pseudotyped with G_K47Q_ or G_R354Q_. Strikingly, retargeted lentiviruses (and in particular those pseudotyped with G_H3-opt-K47Q_ or G_H3-opt-R354Q_) were nearly as efficient as those pseudotyped with G_WT_ (Figure 4C).

### Chimeras specifically target HER2 expressing cells

We then assessed the specificity of the retargeting. For this, we co-cultured HEK293T and HEK293T-HER2^KO^ cells. Each cell line constituted about half of the total cell population at the time of infection by VSVΔG-GFP pseudotyped with different forms of G (Figure 5). VSVΔG-GFP pseudotyped with G_WT_ infected HEK293T and HEK293T-HER2^KO^ cell lines equally efficiently whereas those pseudotyped with G_K47Q_ and G_R354Q_ were virtually unable to infect either cell line (Figure 5). Interestingly, confirming our previous results, lentiviruses pseudotyped with G_H2-opt-K47Q_, G_H2-opt-R354Q_, G_H3-opt-K47Q_, G_H3-opt-R354Q_, G_H4-opt-K47Q_, or G_H4-opt-R354Q_ selectively infected HEK293T cells in the mixed population. A selectivity index can be calculated by dividing the proportion of infected HEK293T cells by the proportion of infected HEK293T-HER2^KO^ cells. The selectivity indexes of the VSVΔG-GFP pseudotyped with the chimeras ranges from 2.23 for G_H1-opt-47Q_ to 5.18 for G_H3-opt-K47Q_ (Figure 5B).

**Figure 5:**
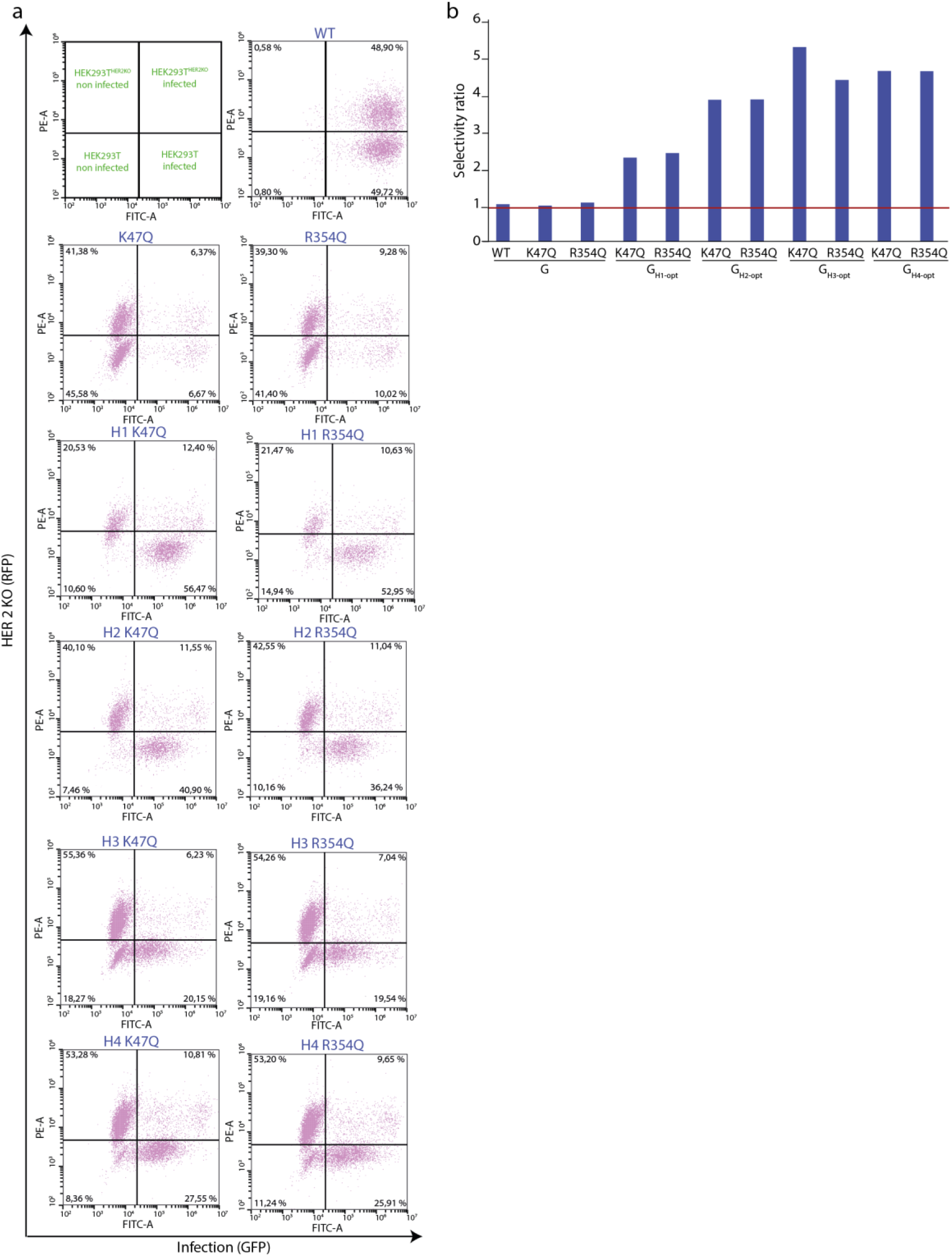
VSV pseudotyped with optimized G_K47Q_ or G_R354Q_ chimeras fused to nanobodies H1 to H4 (recognizing HER2) preferentially infect HEK293T WT cells. **A)** A mix of 50% of HEK293T and 50% of HEK293T-HER2^KO^ (expressing the RFP protein) was infected with VSVΔG-GFP pseudotyped with the indicated G construct. Infected cells were analysed by flow cytometry. Biparametric histograms represent in the x axis the intensity of GFP fluorescence and in the y axis the intensity of RFP. **B)** Selectivity index of VSVΔG-GFP pseudotyped with the chimeras. It was calculated using data shown in panel A) by dividing the proportion of infected HEK293T cells by the proportion of infected HEKT293-HER2^KO^ cells.

## Discussion

One of the major challenges of *in vivo* therapy, including oncovirotherapy and gene therapy, is to target specifically cells of interest. Indeed, the lack of specificity can lead to off-target toxicity but also, in the case of gene therapy (particularly Chimeric Antigen Receptor - CAR-therapies), to trapping of the vector in non-targeted cells. The identification of mutations in VSV G that abolish LDL receptor recognition without affecting fusion activity has opened the possibility of specifically targeting cells of interest.

Indeed, these mutants have been used in combination with a second transmembrane protein, that binds a ligand/receptor of interest, to pseudotype lentivirus, and specifically target them towards a given cell type (28–31). As second transmembrane protein, a dimerized, surface-tethered IL-13 allowed a specific infection of IL-13Rα1-expressing cells (30), while a transmembrane version of an anti-CD19 single-chain antibody fragment (scFv) efficiently infected CD19^+^ B cell lines, but not CD19^−^ T cell lines (30), as well as CD19-expressing HEK293T but not the parental HEK293T cell line (28).

Recently, a gene encoding VSV G, with mutations K47Q and R354Q, fused to a sequence coding for a scFv directed against HER2, was introduced in place of the G_WT_ gene into the genome of a recombinant VSV. The virus was then adapted through serial passages on HER2-expressing mouse mammary tumor line. The resulting virus has a 15-to 25-fold higher titer in HER2-expressing cell lines than in HER2-negative cells (32). Adaptive mutations occured in both G and scFv and therefore are specific to the scFV used in the study.

Here, using experimental evolution, we selected two mutations that optimize G_WT_ backbone to tolerate an amino-terminal fusion with a nanobody recognizing mCherry. The choice of this nanobody (in a context of G still recognizing its natural receptor) had the advantage of limiting a possible GOF which would have allowed the virus to acquire a new receptor (33). Interestingly, these mutations also improved the functionality of most chimeric constructs involving other nanobodies from several libraries. When used to pseudotype VSVΔG-GFP, the optimized chimeric constructs showed a ∼10 fold improvement over the non-optimized chimera. This is particularly interesting given the large number of nanobody libraries available, potentially enabling the targeting of virtually any receptor of interest provided it can induce endocytosis of the viral particle.

Remarkably, the S422I mutation emerged in another context to compensate for the H407A mutation, which abolished the fusion properties of G (19). This mutation stabilizes a β-hairpin structure through hydrophobic stacking of residues I422 between F424 and L430 (19). Here, we show that all hydrophobic residues at this position have the same chimera-optimizing effect suggesting that stabilization of the structural motif is also important in this context. On the other hand, we do not have any molecular explanation for the selection of mutation H22N.

Last but not least, after introducing the K47Q and R354Q mutations in the context of fusion-optimized chimeras with four nanobodies recognizing HER2, we established the proof of concept that it is possible to specifically redirect the infection of pseudotyped VSV or pseudotyped lentiviruses towards this receptor. This specificity is only 5-fold, which may seem limited. However, HEK293T cells do not express much of this receptor on their surface, the specificity could be much greater in the context of a tumor within which cancer cells can express very high levels of this receptor (25). Even using HEK293T cells, our test done using mix population of positive and negative cells showed the capacity or retargeted viruses to specifically infect the subpopulation of positive cells.

Overall, the introduction of the two mutations H22N and S442I (or any other hydrophobic aa in position 422) will therefore make it possible to have functional chimeric glycoproteins both for pseudotyping VSV and lentiviruses, or even for constructing replicative recombinant VSVs, targeting any receptor of interest through the nanobody. These chimera can also be incorporated into the membrane of extracellular vesicles so that they specifically release their cargo into cells of interest for therapeutic purpose (34,35). Of note, it is known that VSV G pseudotyped lentiviruses are poorly effective in infecting certain cell types that do not express LDL-R on their surface (36). The set-up described here may thus also be used in the context of *ex vivo* gene therapy using a mutant chimeric G targeting a receptor of interest different from the LDLR.

## Material and methods

### Cell lines

BSR, a clone of BHK-21 (baby hamster kidney; ATCC CCL-10), and HEK293T (human embryonic kidney expressing simian virus 40 T antigen (SV40T); ATCC CRL-3216) cells were grown in Dulbecco’s modified Eagle’s medium (DMEM) supplemented with 10% fetal calf serum (FCS). SKBR3 (Human Breast Cancer Cell Line; ATCC HTB-30) cells were grown in McCoy’s 5A (Modified) Medium supplemented with 10% FCS.

### Antibodies

Mouse monoclonal antibody directed against VSV G ectodomain was supplied by KeraFAST (8G5F11, dilution 1/1500) and used in cytometry. Mouse anti VSV G (SAB4200695, dilution 1/5000) was purchased from Sigma-Aldrich and used for western blot analysis. Mouse monoclonal 24A1 antibody (kindly provided by Robin Weiss) was used to detect VSV nucleoprotein by immunofluorescence. Mouse anti-HER2 (MA512759, dilution 1/200) was purchased from Invitrogen and used in cytometry. Mouse anti-HER2 (MA513105, dilution 1/200) was purchased from Invitrogen and used in western blot analysis. Goat anti-mouse Alexa fluor 488 (10696113, dilution 1/1000) was purchased from Invitrogen and used as secondary antibody in cytometry. Goat anti-mouse IgG coupled to peroxidase (A4416, dilution 1/5000) was purchased from Merck and used as secondary antibody in western blot analysis.

### Plasmids and cloning

The cDNAs of nanobodies directed against HER2 (H1, H2, H3, H4, H5 and H6) were ordered from Eurofins genomics or Twist Bioscience according to the sequences found on the NCBI database (H1: MP600335, H2: MP600344, H3: MP600340, H4: MP600341, H5: MP600327, H6: MP600331). Chimeric glycoproteins and mutations were created starting from the cloned VSV G gene (Indiana Mudd-Summer strain) in the pCAGGS plasmid (19). For this, overlapping fragments were amplified by PCR and the PCR products and the digested plasmid were assembled using Gibson Assembly kit (New England Biolabs) to obtain the resulting plasmid.

### CRISPR/Cas9-mediated *ERBB2* gene knockout

*ERBB2* gene (encoding HER2 protein) was knocked out using CRISPR/Cas9 KO and HDR (Homologous directed recombination) plasmids from Santa Cruz biotechnology (37). Briefly, HEK293T were transfected with 1µg of a pool of three plasmids [CRISPR/Cas9 KO ErbB2/HER2 (h2) (sc-400138-KO-2)] each encoding Cas9 nuclease but distinct ERBB2-specific guide RNA and 1µg of plasmid HDR [ErbB2/HER2 (h2) (sc-400138-HDR-2)] used for selection. HEK293T-HER2^KO^ cells were then selected using 0.75µg/ml of puromycin. The invalidation of ERBB2 was confirmed by cytometry and western blot.

### Cell surface expression

To quantify the expression of G protein on the cell surface, HEK293T cells plated on six-well dishes at 70% confluence were transfected with G-expressing plasmid using the calcium phosphate method. At 24 h after transfection, cells were detached with 1ml of 2% FCS in PBS (phosphate buffer saline, 10 mM Na_2_HPO_4_ , 1.8 mM KH_2_PO_4_ , 137 mM NaCl, 2,7 mM KCl, pH 7.4), 200µl were harvested in a 96-well plate, followed by centrifugation at 350 g for 3 min. Cells were then incubated with a 1:1,500 dilution of mouse monoclonal anti-G antibody (8G5F11) in PBS at 4 °C for 30min. Cells were washed twice in PBS, incubated with a 1:1000 dilution of goat anti-mouse Alexa Fluor 488 (Invitrogen) at 4°C for 30 min., and rinsed in PBS. After resuspension in 100 μl of 2% FBS-PBS, the fluorescence of 10,000 cells from each population was determined by flow cytometry using a Cytoflex S cytometer. The median fluorescence intensity (MFI) of the transfected cells expressing G was quantified by flow cytometry. The relative cell surface expression of transfected cells was determined as follows: (MFI for the mutant)/(MFI for the WT). For each mutant, the percentage given is the average of at least three independent experiments.

### Generation of rVSV-G_C11_ recombinant

Plasmid pVSV-FL(+) expressing the 11,161-nucleotide (nt) positive-strand full-length VSV RNA sequence and plasmids pBS-N, pBS-P, and pBS-L, respectively, encoding N, P, and L proteins were kindly provided by John K. Rose (Yale University, New Haven, CT, USA). The chimeric G gene of the VSV Indiana serotype (Mudd-Summers strain) in fusion with that encoding the nanobody C11 was inserted into the original full-length genomic plasmid pVSV-FL(+) (38) using two unique restriction sites in pVSVFL(+), MluI in the 5′ noncoding sequence of the glycoprotein (G) gene and NheI present in a sequence introduced between G and L genes, after removal of the corresponding VSV Indiana Mudd-Summers G gene. Recombinant rVSV-G_C11_ was recovered in BSR cells as described (39). Briefly, BSR cells were infected with recombinant vaccinia virus expressing bacteriophage T7 RNA polymerase (vTF7-3) at a multiplicity of infection of 10. After 1 h, BSR cells were co-transfected with the pVSV-FL(+), pBS-N, pBS-P, and pBS-L plasmids using Lipofectamine 2000 (Invitrogen) in the presence of 10 μg/ml 1-β-d-arabinofuranosylcytosine (araC; Sigma). The recovery of the recombinant was supported by the expression of functional WT VSV G protein in *trans* from a plasmid (pcDNA3.1 G WT) which was cotransfected with plasmids pVSV-FL(+), pBS-N, pBS-P, and pBS-L.

### Experimental evolution of rVSV-G_C11_ and sequencing

Experimental evolution rVSV-G_C11_ was achieved by serially passing virus at very low MOI (∼3 10^−4^) on BSR cells. At each passage, the virus was titrated by plaque assay. Infected cells were lysed 6 h post infection in RNAnow (Ozyme). RNA were extracted and mRNA were reverse transcribed using a T_12_CAT primer and the cDNA corresponding to M and G mRNAs were amplified by PCR. The PCR fragments were sequence by the Sanger’s method (Eurofins).

### Virus purification

rVSV-G_C11_ was propagated on BSR cells. Sixteen hours post-infection, cell debris were eliminated from the supernatant by filtration (0.22 µm). The virus was then pelleted by centrifugation at 4°C for 45 min at 25,000 rpm in a sw28 rotor (Beckman) and gently suspended in dilution buffer (150 mM NaCl, 10 mM Tris-HCl pH 7.5, 2 mM EDTA).

### Preparation of VSVΔG-GFP pseudotypes

HEK293T cells at 80% confluence were transfected by pCAGGS encoding WT or chimeric Gs using the calcium phosphate method. Twenty-four hours after transfection, cells were infected with VSVΔG-GFP pseudotyped with VSV G_WT_ at an MOI of 3. One hour post infection, 1ml of complete DMEM (*i.e.* supplemented with 10% FCS and Penicillin/Streptomycin) was added. At two hours post infection (p.i.), cells were washed to remove residual viruses from the inoculum and 2ml of complete DMEM were added. Cell supernatants containing the pseudotyped viral particles were collected at 16 h p.i. and clarified by centrifugation at 4°C, 600 g, 15 min.

### Titration of VSVΔG-GFP pseudotypes

The infectious titers of pseudotyped VSVΔG-GFP were calculated by counting cells expressing the GFP using a flow cytometer at 16 h p.i. and using the equation MOI=−ln[p(0)] where MOI is the multiplicity of infection and p(0) is the proportion of non-infected cells.

### Preparation of lentivirus pseudotypes

HEK-293T cells at 80% confluence were transfected by pCAGGS encoding G_WT_ or chimeric Gs, psPAX2 encoding HIV-1 gag and pol (12260 purchased from Addgene), and pHIV-eGFP encoding eGFP (21373 purchased from Addgene) using polyethylenimine (PEIMax; Polyscience) at 1mg/ml. At 18h after transfection, media was replaced with fresh complete DMEM. After 28 h, cell supernatants containing the pseudotyped viral particles were collected. Supernatants were clarified by centrifugation at 4°C, 600 g, 15 min.

### Titration of lentivirus pseudotypes

The infectious titers of the pseudotyped lentiviruses were calculated by counting cells expressing the GFP using a flow cytometer at 48 h post transduction and using the equation MOI=−ln[p(0)] where MOI is the multiplicity of infection and p(0) is the proportion of non-infected cells.

### Western blot analysis

Supernatant containing VSVΔG-GFP pseudotypes were harvested and centrifugated at 2500 rpm during 15 min. at 4°C. Clarified supernatant were concentrated by ultracentrifugation at 40,000 g during 45 min. at 12°C. After resuspension of the pellets in 100 µl of TNE (20 mM Tris pH 7.5, 50 mM NaCl, 2 mM EDTA) buffer, 60 µl were harvested and after the addition of 20 µl of Laemmli Buffer, pseudotyped viruses were lysed at 95°C.

HEK293T, HEK293T-HER2^KO^ and SKBR3 cells were centrifugated at 600 g for 10 min. at 4°C. Pellets were resuspended using RIPA buffer (Tris-HCL 25 mM pH 8.8, NaCl 50 mM, 0.5% Nonidet P-40, 0.1% SDS) and a cocktail of protease inhibitors (Roche Diagnostic, #11-836-170-001) and incubated on ice for 30 min. before 2 min. of centrifugation at 9500 g. Lysed extracts were denatured in Laemmli Buffer at 95°C.

For the western blot, proteins were separated by electrophoresis on 12% SDS-PAGE and electro-transferred into a nitrocellulose membrane before blocking of the membrane with 5% of skimmed milk in Tris Buffer Saline (TBS, pH 7.5 #ET220-B, Euromedex), supplemented with 0,1% Tween 20, for 45min at room temperature. The membranes were then incubated overnight at 4°C with the corresponding primary antibodies in TBS, 0,1% Tween20, 5% skimmed milk. After the incubation, membranes were rinsed three times with TBS 0,1% Tween20 and incubated with the secondary antibody for 1 h at room temperature. After washing, the membranes were scanned with the Odyssey infrared imaging system (LI-COR, Lincoln, NE)

### Electron microscopy

Samples of purified virions diluted in 150 mM NaCl, 50 mM Tris-HCl, pH 7.5, or dialyzed against a buffer containing150 mM NaCl and 50 mM MES at pH pH 5.5, were analyzed by conventional electron microscopy using the negative staining method. Three microliters of sample suspension were deposited on an air glow-discharged 400 mesh copper carbon-coated grid for 1 minutes. The excess of liquid was blotted, and the grid stained with sodium phosphotungstic acid adjusted to pH (7.5 or 5.5). The grids were visualized at 100 kV with a Tecnai 12 Spirit transmission electron microscope (Thermo Fisher, New York NY, USA) equipped with a K2 Base 4k x 4k camera (Gatan, Pleasanton CA, USA.

### Statistical analysis

Statistical analysis were performed using Prism (GraphPad software). Data comparison was performed using Student t-test. Statistical details of experiments are indicated in the figure legends. Results are indicated as mean ± SD.

## Acknowledgements

This work was supported by a grant from a grant from Fondation pour la Recherche Médicale (FRM), France, to YG (EQU202103012746), a grant from l’Agence pour la Recherche sur le Cancer (ARC), France, to AAA and by a grant from la Banque Publique d’Investissement (Projet ETINCELL) including a salary for ID. The work in FP lab work was supported by the CNRS and by the Institut Curie as well as by Labex Cell(n)Scale (ANR-10-LBX-0038 part of the IDEX PSL no. ANR-10-IDEX-0001-02. The authors would like to thank Sandrine Moutel from the Antibody facility of the Institute Curie for the kind gift of specific nanobody sequences.

